# The toxic nature of murine amylin and the immune responsivity of pancreatic islet to conformational antibody in mice

**DOI:** 10.1101/175539

**Authors:** Luiza C. S. Erthal, Luana Jotha-Mattos, Flávio Alves Lara, Sabrina Alves dos Reis, Bernardo Miguel de Oliveira Pascarelli, Cinthia Melo Costa, Kleber L. A. Souza, Luís Maurício T. R. Lima

## Abstract

The human amylin is a pancreatic peptide hormone cosecreted with amylin and found in hyperhormonemic state along with insulin in subclinical diabetes. Amylin has been associated with the pathology of type 2 diabetes, particularly due to its ability to assembly into toxic oligomers and amyloid speciments. On the other hand, some variants such as murine amylin has been described as non amyloidogenic, either *in vitro* or *in vivo*. Recent data have demonstrated the amyloid propensity of murine amylin and the therapeutic analogue pramlintide, suggesting a universality for amylin amyloidosis. Here we report the amyloidogenesis of murine amylin, which showed lower responsivity to the fluorescent probe thioflavin T compared to human amylin, but presented highly organized fibrilar amyloid material. The aggregation of murine amylin also resulted in the formation of cytotoxic specimens, as evaluated *in vitro* in INS-1 cells. The aggregation product from murine amylin was responsive to a specific antibody raised against amyloid oligomers, the A11 oligomer antibody. Pancreatic islets of swiss mice have also shown responsivity for the anti-oligomer, indicating the natural abundance of such specimen in rodents. These data provide for the first time evidences for the toxic nature of oligomeric assemblies of murine amylin and its existence in non-transgenic mice.

**Highlights:**

- Murine amylin forms oligomer species and amyloid fibrils *in vitro*
- The murine amylin aggregation product display cellular toxicity
- A11 anti-oligomer antibody recognizes murine amylin *in vitro*
- Non-transgenic mice display immunoresposivity to anti-oligomer in pancreatic islet

## Introduction

The existence of amyloid deposits in pancreas has been known for over a century [1]. More recently, the nature of these depots in pancreas of diabetic human and cats were confirmed to be amylin aggregates [2, 3]. The lack of detectable amyloid material in some other animals possessing proline in specific positions of their amylin [4] raised a possible role of this imino acid in hampering the formation of amyloid assemblies, most likely due to the beta-disrupting character of prolines [5]. Such discovery prompted the development of the triple proline (Pro^25,28^,^29^) human amylin analogue known as pramlintide, which entered the pharmaceutical market in 2005 [6]. Formulated in acetate pH 4.0, pramlintide displays chemical and physico-chemical stability, although the intrinsic propensity for fast amyloid aggregation is still dominant in higher pH [7]. Murine amylin is also prone to amyloid aggregation *in vitro* [8, 9], which collectively suggest an universal feature of amylin analogues.

The amylin analogue from pufferfish has recently been described as capable to aggregate into amyloid material although not detectable by thioflavin T (ThT) [10]. In conjunction these data indicate that despite the propensity of these amylin analogues to assembly into amyloid material, quantitative detection might have been an historical issue, both *in vitro* and in biological specimens. In fact, previous studies have also reported amyloid-related properties for murine amylin *in vitro* [11–13]. Some other *ex vivo* studies have also detected oligomer-like immunoreactive material (OLIM) [14] and congo-red birefringent fibers [15] in the endocrine pancreas of rodents, though considered of minor importance.

We raised the hypotheses weather it would exist a universal propensity for the oligomeric assembly and amyloid formation of amylin analogues, whose detection would be hampered both by ligands such as buffers as observed for murine amylin [9] and pramlintide [7], among other technical details. In this work we report the investigation of the toxic nature of murine amylin and the natural abundance of oligomer-like immunoreactive material in pancreatic islets of non-trangenic mice.

## MATERIAL AND METHODS

### Materials

High-purity (>95 %) murine (CAS 124447-81-0) and human (CAS 122384-88-7) amylin, was obtained from Genemed Biotech Inc (CA, USA). Stock amylin solutions were prepared at 10 mg/mL (2.55 mM) in DMSO and stored at -20 °C. All other reagents were of analytical grade.

### Amylin aggregation assay

Amylin amyloid aggregation was conducted at 25 °C in a 96-well flat-bottom black plate (Corning cat #3915) sealed with Crystal Clear Tape (Duck Brand, Avon, OH), containing 50 μM amylin (from stock at 2.55 mM in DMSO), 20 μM ThT (from 3 mM stock in water), 50 mM Na_2_HPO_4_ pH 7.4. Measurements were performed at 3 min time interval with 3 sec aggitation prior top-read fluorescence acquisition with excitating and emission set at 440 nm and 482 nm respectivelly, and cut-off filter 475 nm. The aggregation kinetics were analysed with the empirical four-parameters sigmoid function with Sigma Plot 12.5 (Systat Software, Inc., CA, USA) as previously described [7, 9].

### Transmission electron microscopy (TEM)

Aliquots of 5 μL from the aggregation kinetic assays were applied to 300 mesh, Formvar-coated Cu grids, the excess removed and the grid was subjected to negative stain with 2% uranyl acetate, pH 4.8, for about 30 s protected from light, the excess removed and the grid was air-dried. The stained samples were observed in a transmission electron microscope (Tecnai G2, FEI; DIMAV-INMETRO) operating at 80 to 120 kV.

Immunochemistry was performed at room temperature by applying aliquots aggregation kinetic assays onto 300 mesh, Formvar-coated Cu grids, the excess removed, blocked with blocking buffer (927-40000 Odyssey® Blocking Buffer, PBS-based), treated with primary antibodies OC (anti amyloid fibril-like material; Cat #AB2286, Merck-Millipore) or A11 (anti prefibrillar oligomer-like material; Cat #AB9234, Merck-Millipore), washed 2 times with blocking buffer, treated with secondary antibody immunogold (10 nm particles) and washed for 6 times, and stained with uranyl acetate. The stained samples were observed in a transmission electron microscope (Tecnai G2, FEI) operating at 80 to 120 kV.

### Immunodetection of prefibrillar oligomer-like immunoreactive material (OLIM) and fiber-like immunoreactive material (FLIM)

The detection of OLIM and FLIM in samples from the *in vitro* aggregation assays of amylin (above) was performed by dot blot. In brief, the PVDF membrane (IPFL00010 | Immobilon-FL PVDF – Merck-Millipore) was incubated for 1 minute in 100% methanol. The membrane and three filter papers were then placed in transfer buffer (20% methanol and running buffer (25 mM Tris, 192 mM Glycine pH 8.3) and assembled in the dot blotting system Bio-Dot® SF Microfiltration Apparatus 170-6542 (BioRad, Brazil). 100 μL of each amylin sample from aggregation assay was applied in each slot and the dot blotting system was placed in a vacuum pump to draw samples through the membrane. The system was washed with distilled water once and the membrane was incubated with 5 mL of blocking buffer (927-40000 Odyssey® Blocking Buffer – PBS, LI-Cor® Biotech) at 4 °C for 2h followed by incubation overnight at 4 °C with the primary antibody OC (anti amyloid fibril-like material; Cat #AB2286, Lot #NG1865063, Merck-Millipore) or the antibody A11 (anti prefibrillar oligomer-like material; Cat #AB9234, Lot #2387440, Merck-Millipore) at a dilution of 1:1,000. The primary antibody was then removed and the membrane was washed out for four times with PBS in the presence of 0.1% Tween 20 for 5 minutes each round. The secondary antibody (IRDye 800CW; Li-Cor Biosciences) at a dilution of 1:10,000 was then incubated for 1 hour at 4 °C under protection of light. The blotting was revealed in Odyssey Infrared Imaging System (Li-Cor Biosciences; CENABIO-UFRJ) using the intensity set #5 for human amylin and the set #7 for murine amylin (due to its lower signal compared to results performed in parallel with the more immune responsive human amylin).

### Immunohistochemistry

Swiss male mice (CEMIB-UNICAMP-Campinas) were housed in a mini-isolators in ventilated racks (VentiLife 112, Alesco, Brazil) within a temperature-controlled room with a light-dark cycle of 12 h. Water and standard chow diet (Biobase, Cat #9301, Brazil) were available ad libitum. 12 weeks-old mice were fasted for 4 h and euthanized by decaptation. Pancreatic tissues were collected and fixed in 4 % paraformaldehyde, embedded in paraffin and cut in 5 μm sections for further immunohistochemistry with anti-murine amylin (in-house derived, from rabbits, produced by EJ-APC) and A11 (anti prefibrillar oligomer-like material; Cat #AB9234, Merck-Millipore). Alexa633-conjugated α-rabbit (1:400, Invitrogen, USA) was employed as secondary antibody. Unspecific binding was monitored using as primary antibody non-immune rabbit serum. Paraffin sections were dewaxed in xylol, hydrated in graded ethanol and distilled water. The sections were then heated in 10 mM sodium citrate buffer (pH 6.0) for 10 min in 700W microwave oven followed by 30 min to cool down to room temperature to retrieve the antigens. After blocking with PBS solution containing 5% bovine serum albumin, 0.25% Triton X-100 and 10% normal goat serum, sections were incubated overnight in a humid chamber (4°C) with the respective primary antibody, followed by 3 washes (5 min each), and then incubated with secondary antibodies diluted in the same blockage solution for 90 min at room temperature. After a final series of washes in PBS (3 x 5 min each), cell nuclei were counterstained with DAPI (4’,6’-diamidino-2-phenylindole, Sigma-Aldrich) and each slide received a coverslip containing a drop of SlowFade® antifade solution (Molecular Probes, OR, USA). Images were performed during the analysis of three biological replicates using a Zeiss AxioObserver Z1 fluorescence microscope (Carl Zeiss, Heidenheim, Germany), using a 630 nm LED (Colibri Iluination System) as excitation source with a Zeiss 50 Filter set. Acquisition was performed using a HMR CCD controlled by software Axiovision (Carl Zeiss, Heidenheim, Germany). Based in the signal/noise ratio between immunostained and negative controls, a threshold was established and positive cells were quantified by direct counting. This protocol was approved by the Institutional Bioethics Committee on Animal Care and Experimentation at UFRJ (# FARMACIA05/2012).

### Toxicity assay in cell-cultures of INS-1

Insulin-secreting INS-1 cells were cultured in RPMI medium, supplemented with 10 mM glucose, 10 % (v/v) fetal calf serum (FCS), 1 mmol/L sodium pyruvate, 50 μmol/L 2-mercaptoethanol, 2 mmol/L glutamine, 5 mmol/L of HEPES, penicillin, and streptomycin in a humidified atmosphere at 37 °C and 5% CO_2_. For MTT assays, 96-well flat-bottom plates were used. Cells were platted at an initial density of 7.5 x 104 (experiments of 24h incubation) or 5 x 104 (48h incubation) per well and allowed to attach for 15h. Thereafter cells were exposed to amylin at 50 μM from the aggregation assays. Control assays were INS-1 cells and vehicle only, in the absence of amylin. The cellular viability was accessed after incubation the period using a microplate based MTT assay [16]. Viability was expressed in % of the MTT absorbance at 550 nm in the absence of amylin.

## RESULTS

### Kinetics of amylin aggregation

We have conducted an amyloid aggregation assay of both human and murine amylin in phosphate buffer. Both variants show a typical aggregation profile characterized by a lag phase followed by an exponential increase in ThT fluorescence over time converging to a plateau (**Fig. 1**). Human amylin reaches the plateau within about 10 h, while no signal of murine amylin ThT binding is detected for up to 27 h, until a rise in ThT fluorescence is observed, reaching about 20 % of the magnitude of the human amylin signal under similar aggregation conditions. The dissimilar magnitude of the total ThT signal may be due to the total amount of ThT bound per total amyloid material, the quantum yield and / or the total amount of amyloid material formed. Besides, these data evidences that both human and murine amylin show a typical kinetic profile of amyloid aggregation as monitored by ThT and that human amylin show a faster aggregation profile compared to murine amylin. Further possible limitation in amyloid aggregation assays of murine amylin may be due to the use of tris buffer, which hampers amylin aggregation [7, 9].

**Figure 1.**
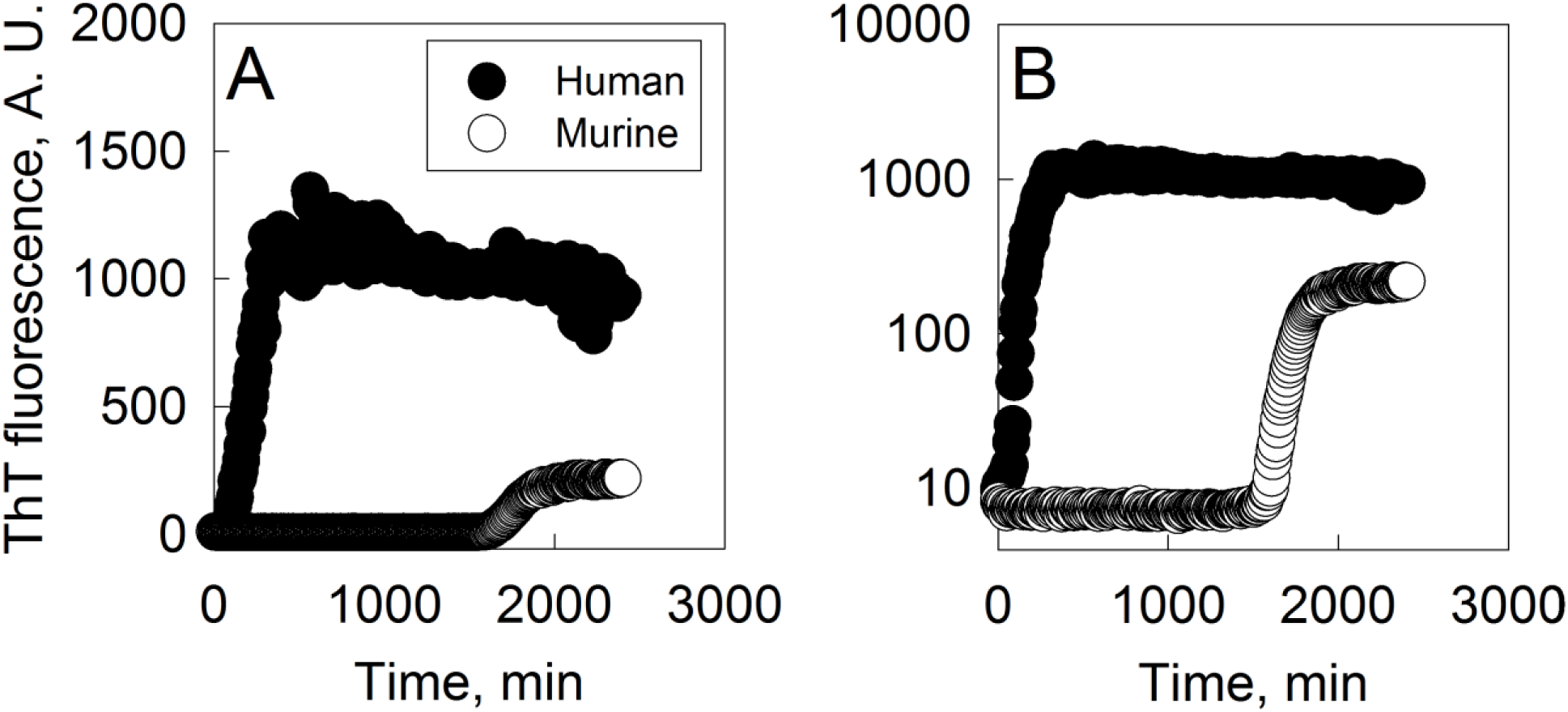
Kinetics of amyloid aggregation of human and murine amylin. Amylin (50 μM) was incubated at 25 °C in 50 mM phosphate buffer pH 7.4 and 20 μM ThT and the aggregation was followed by fluorescence (ex 440 nm, em 482 nm, filter 475 nm) with 3 sec. agitation prior to measurements with 3 min interval. ThT fluorescence axis: A) linear scale; B) log scale.

### Morphological and Immunochemical properties of human and murine amylin aggregates

In order to get further insight on the nature of the aggregate material formed by human and murine amylin, we performed a immunoblot assay by using the A11, an antibody sensible to amylin prefibrillar oligomer-like immunoreactive material (OLIM), and the OC, an antibody sensible to amyloid fiber-like immunoreactive material (FLIM) [17–19]. The products of amylin aggregation assays were adhered onto Formvar films and observed by transmission electron microscopy (TEM). For both human and murine amylin, mature fibrils and oligomers were observed by transmission electron microscopy (TEM), for both 3 days and 7 days incubation (**Fig. 2**). We have found long fibrils mounting twisted fibers, characteristic of amyloid material [20] in both human and murine amylin (**Fig. 2**), confirming the existence of amyloid material in the aggregated samples. While both the human and murine amylin aggregates showed response for the A11 (anti-OLIM antibody) at short incubation period, the immunoblot assay revealed OC (anti-FLIM antibody) responsivity at about day 3 of incubation for both human and murine amylin (**Fig. 3)**. These aggregates were not detectable in both human and murine amylin directly tested from soluble peptide stock solution in DMSO (not shown). Under TEM we could observe the specificity of these antibodies, able to recognize small material assembly and no binding to mature fiber (**Fig. 4**).

**Figure 2.**
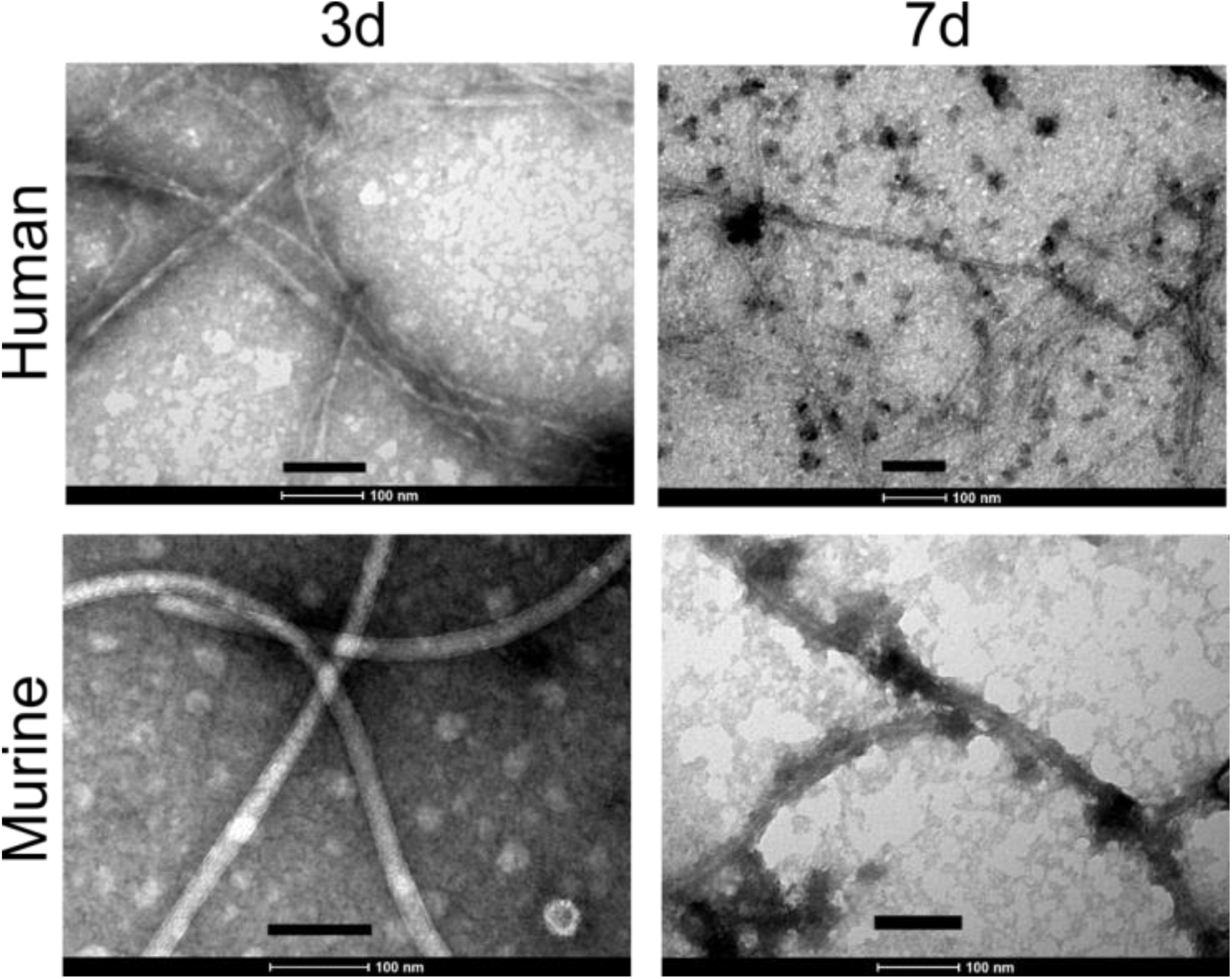
Morphological characterization of the amylin aggregates. Murine and human amylin (50 μM) were incubated at 37 °C in 10 mM Na_2_HPO_4_ pH 7.4 and at given time interval the material were evaluated by TEM. Bar scale = 100 nm.

**Figure 3.**
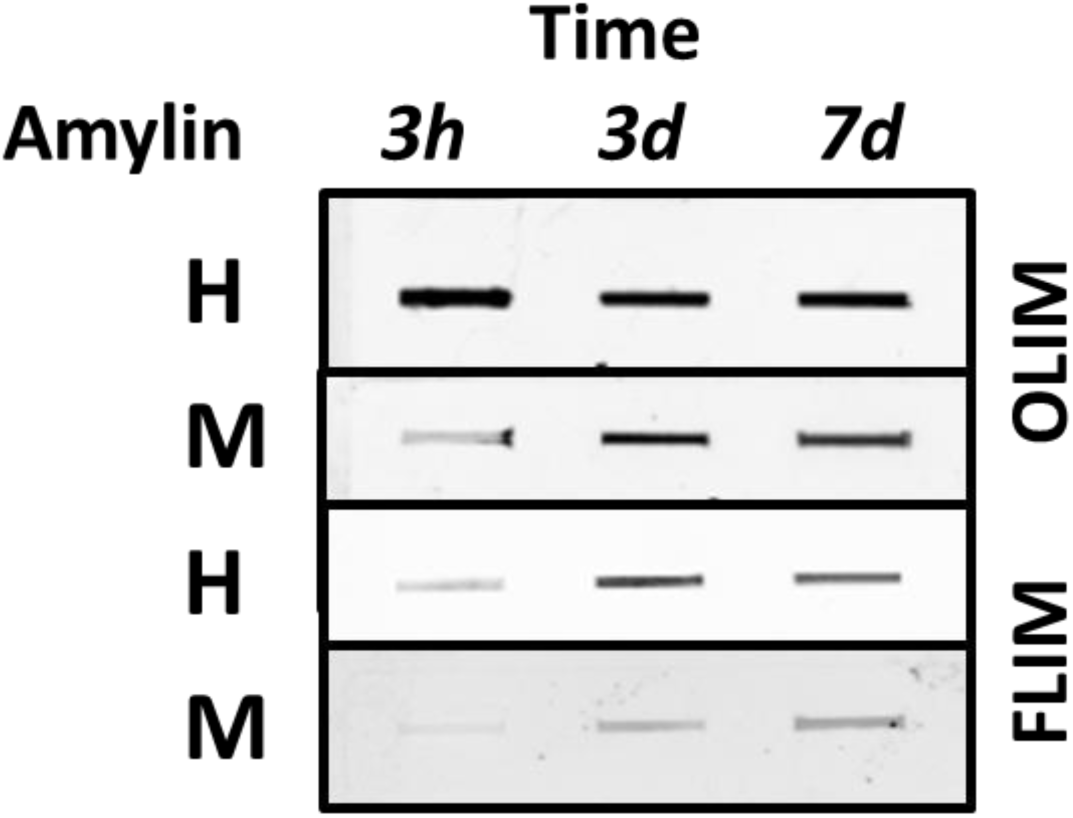
Conformacional immunogenicity of the amylin aggregates. Human (H) and murine (M) amylin (50 μM) were separately incubated in 10 mM Na_2_HPO_4_ pH 7.4 at 37 °C and the formation of oligomer-like immunoreactive material (OLIM) and amyloid fiber-like immunoreactive material (FLIM) were accessed by dot-blot analysis.

**Figure 4.**
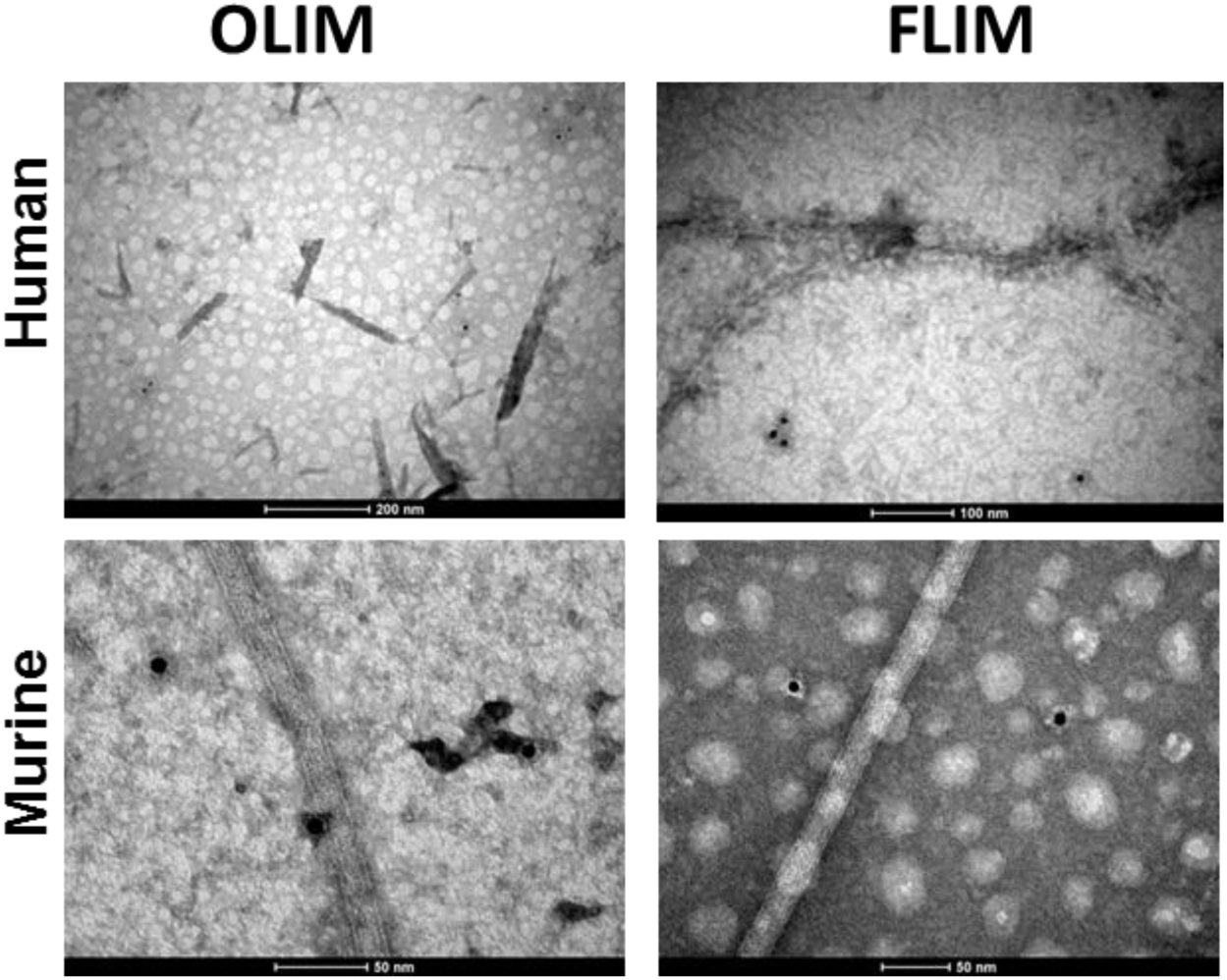
Immunochemical characterization of amyloid aggregates. Human (H) and murine (M) amylin (50 μM) were separately incubated in 10 mM Na_2_HPO_4_ pH 7.4 at 37 °C and the tested for detection by oligomer-like immunoreactive material (OLIM) and amyloid fiber-like immunoreactive material (FLIM) by with further binding of secondary immunogold (10 nm size).

During dot-blot data acquisition, we observed that the signal for both antibodies in response to murine amylin samples were lower than those used for human amylin, requiring different instrumental setting to compensate for these features. In fact, the amyloid aggregates of murine amylin seem less responsive to ThT [8, 9]. We reasoned whether the typical lack of signal for oligomers in rodents could be an issue of tinctorial responsiveness or a dissimilar antibody titer between human and murine amylin aggregates, a feature also observed for other amylin variants [10]. In fact, such as OLIM could be detected in pancreas of elder humans without diagnosed overt diabetes [21], although a hyperhormonemic subclinical diabetes condition [22, 23] could be responsible for such outcome. We were able to detect OLIM responsiveness for the A11 in 16-week old mice (**Fig. 5 and Supp. Fig. 1 and Supp Fig. 2**). These data indicate the responsivity of both human and murine amylin assemblies for the immunodetection by the A11 (anti-OLIM) both *in vitro* and in biological specimens.

**Figure 5.**
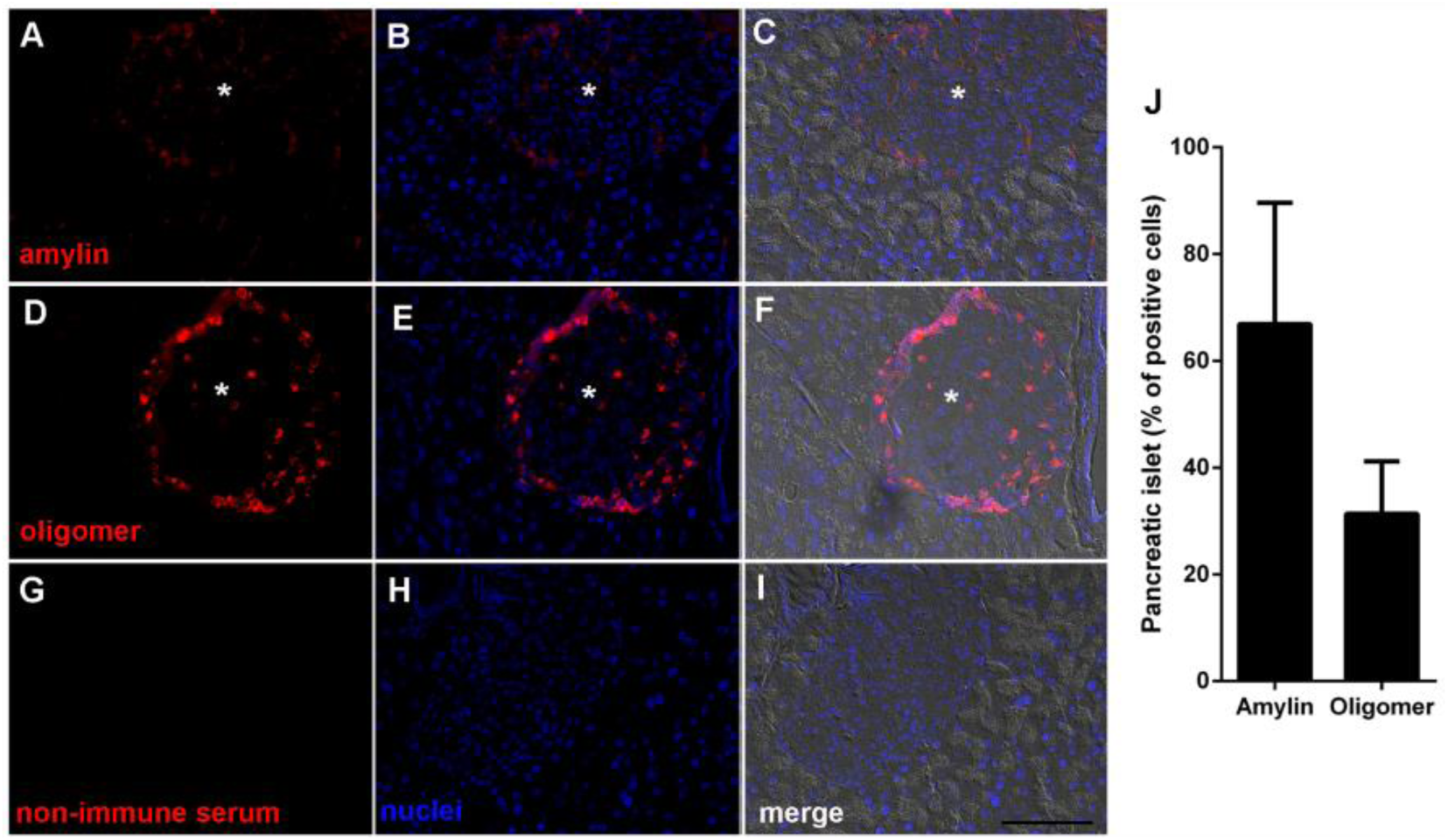
Pancreatic islet oligomers in mice. 12 weeks-old mice pancreas were immunostained by primary antibodies against amylin or soluble oligomers as indicated. Nuclei were stained in blue by DAPI. The staining for amylin and oligomer is observed within the islet (asterisks) as expected. Control images were performed with consecutive section, omitting the primary antibodies (anti-amylin or anti-oligomer) and using a non-immune serum instead. Images are representative of 100 fields from three different animals. Scale bar means 50μm. Islets cells presenting positive signal for amylin or oligomer were quantified. Additional images can be found in **Supp. Fig. 1 and Supp. Fig. 2**.

### Cellular toxicity of human and murine amylin

The toxicity of the amylin aggregates was evaluated in insulinome INS-1 cell line by monitoring the MTT reduction assay. Both human and murine amylin were allowed to aggregate for varying time intervals in phosphate buffer at pH 7.4 and further assayed for toxicity. We observed a reduction in the viability of cell as accessed by the MTT assay (**Fig. 6**), both for human and murine amylin. These data indicate that the murine amylin aggregates contain material displaying toxic behavior for the insulinome cell lines, a similar behavior to that observed for human amylin aggregates. Although some authors suggest that oligomers are toxic specimens [12, 17, 24, 25], for the present moment one cannot assure which fraction of the amyloid aggregates generated in the present study are more likely to behave as toxic agents. Further investigation will require a careful fractioning and extensive structural and toxicological characterization of the material.

**Figure 6.**
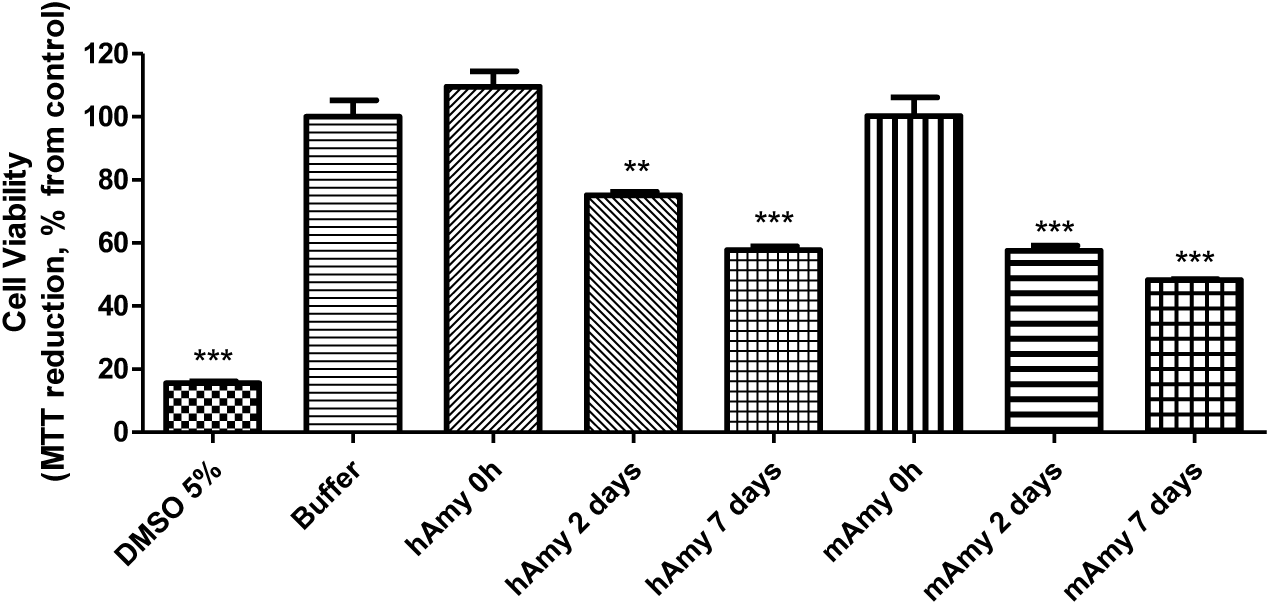
Toxicity assay in INS-1 insulinome cell line. Amylin (50 μM) was allowed to aggregate in 10 mM Na_2_HPO_4_ pH 7.4 at 37 °C and assayed for cell toxicity by MTT assay. ** P<0.001 and *** P<0.0001, compared to control (buffer only).

## DISCUSSION

We have shown here that the product of both human and murine amylin aggregation assay *in vitro* contain both amyloid fibrils and species reactive to anti-oligomer antibody, and that they present similar toxic pattern to INS-1 insulinome cell line. Moreover, our data demonstrate that OLIM antibody is able to selectively recognize and binding to the islet of non-transgenic swiss mice, suggesting the existence of oligomer-like immunoreactive material in wild-type murine pancreas.

The historical lack of observation of amyloid material in murine models has been interpreted as a lack of the amyloidogenic pathway in these organisms, which has been sustained by the inability of detection of ThT response in aggregation process *in vitro.* However, recent data have shown that ThT is not an absolute probe for amyloid material, given that murine amylin [8, 9] and puffer-fish amylin [10] and even the more stable amylin analogue pramlintide [7] can form amyloid material, with ThT responsiveness at lower threshold for detection when compared to the product of human amylin aggregation assay.

For over 25 years, the misguided lack of observation of amyloid aggregates both *in vitro* and *in vivo* has delineated the conventional wisdom that proline-rich amylin variants do not evolve to amyloid aggregates (Westermark *et al.* 1990). Indeed, several independent groups have reported their lack of detection of amyloid deposits in rodents, prompting for the development of transgenic models overexpressing human amylin. However, as from an analytical perspective, absence of signal is not equal to absence of matter. Classical evaluation of amyloid deposits involved tintorial detection by histochemical techniques [1, 26, 27], which show low sensitivity to amyloid material from murine amylin [8]. More recently, antibodies anti-oligomer-like and anti-fibril-like immunoreponsive material (OLIM and FLIM, respectivelly) have been developed and tested against a variety of amyloid polypeptides, showing adequate responsiveness [28][29].

Our group have shown for the first time that murine amylin does form low and high order oligomers and amyloid fibers *in vitro* (Palmieri *et al.* 2013b). Although the morphologic characteristics of these murine aggregates show typical signatures of amyloid material such as x-ray diffraction pattern, fibrils on transmission electron microscopy and atomic force microscopy [8, 9], the tintorial properties of these aggregates were less pronounced when compared to the material obtained from human amylin, limiting the detection of amyloid material from murine amylin. However, we have shown here for the first time to the best of our knowledge that the classical A11 and OC antibodies against OLIM and FLIM are responsive to murine amylin aggregate material, which could be used to specifically detect OLIM in pancreatic islets of non-transgenic mice. These results indicate the potential of murine amylin populating the oligomers in natural abundance *in vivo,* and the use of immunochemistry in the investigation of such events in non-transgenic murine animals models of amyloid-related pathologies.

## Acknowledgments

We would like to thank Dr. Ivone Rosa for excellent technical services, to Dr. Guilherme Augusto (IBqM) for helpful assistance in setting-up dot-blot assays, and to DIMAV-INMETRO and CENABIO-UFRJ for providing access to their analytical platforms. This work was supported by the Coordenação de Aperfeiçoamento de Pessoal de Nível Superior (CAPES), Conselho Nacional de Desenvolvimento Científico e Tecnológico (CNPq), Fundação de Amparo à Pesquisa do Estado do Rio de Janeiro Carlos Chagas Filho (FAPERJ). The funding agencies had no role in the study design, data collection and analysis, or decision to publish or prepare of the manuscript.

## Ethical approval

“All applicable international, national, and/or institutional guidelines for the care and use of animals were followed. All procedures performed in studies involving animals were in accordance with the ethical standards of the institution or practice at which the studies were conducted.”

## REFERENCES

1. Opie EL (1901) The relation oe diabetes mellitus to lesions of the pancreas. Hyaline degeneration of the islands oe langerhans. J Exp Med 5:527–540.

2. Cooper GJ, Willis AC, Clark A, et al (1987) Purification and characterization of a peptide from amyloid-rich pancreases of type 2 diabetic patients. Proc Natl Acad Sci U S A 84:8628–8632.

3. Westermark P, Wernstedt C, Wilander E, et al (1987) Amyloid fibrils in human insulinoma and islets of Langerhans of the diabetic cat are derived from a neuropeptide-like protein also present in normal islet cells. Proc Natl Acad Sci U S A 84:3881–3885.

4. Westermark P, Engström U, Johnson KH, et al (1990) Islet amyloid polypeptide: pinpointing amino acid residues linked to amyloid fibril formation. Proc Natl Acad Sci U S A 87:5036–5040.

5. Deber CM, Therien AG (2002) Putting the β-breaks on membrane protein misfolding. Nat Struct Mol Biol 9:318–319. doi:10.1038/nsb0502-318

6. McQueen J (2005) Pramlintide acetate. Am J Health-Syst Pharm AJHP Off J Am Soc Health-Syst Pharm 62:2363–2372. doi:10.2146/ajhp050341

7. da Silva DC, Fontes GN, Erthal LCS, Lima LMTR (2016) Amyloidogenesis of the amylin analogue pramlintide. Biophys Chem 219:1–8. doi:10.1016/j.bpc.2016.09.007

8. Palmieri LC, Melo-Ferreira B, Braga CA, et al (2013) Stepwise oligomerization of murine amylin and assembly of amyloid fibrils. Biophys Chem 180–181:135–144. doi:10.1016/j.bpc.2013.07.013

9. Erthal LCS, Marques AF, Almeida FCL, et al (2016) Regulation of the assembly and amyloid aggregation of murine amylin by zinc. Biophys Chem 218:58–70. doi:10.1016/j.bpc.2016.09.008

10. Wong AG, Wu C, Hannaberry E, et al (2016) Analysis of the Amyloidogenic Potential of Pufferfish (Takifugu rubripes) Islet Amyloid Polypeptide Highlights the Limitations of Thioflavin-T Assays and the Difficulties in Defining Amyloidogenicity. Biochemistry (Mosc) 55:510–518. doi:10.1021/acs.biochem.5b01107

11. Green J, Goldsbury C, Mini T, et al (2003) Full-length rat amylin forms fibrils following substitution of single residues from human amylin. J Mol Biol 326:1147–1156.

12. Milton NGN, Harris JR (2013) Fibril formation and toxicity of the non-amyloidogenic rat amylin peptide. Micron Oxf Engl 1993 44:246–253. doi:10.1016/j.micron.2012.07.001

13. Lopes DHJ, Colin C, Degaki TL, et al (2004) Amyloidogenicity and cytotoxicity of recombinant mature human islet amyloid polypeptide (rhIAPP). J Biol Chem 279:42803–42810. doi:10.1074/jbc.M406108200

14. Gurlo T, Ryazantsev S, Huang C, et al (2010) Evidence for proteotoxicity in beta cells in type 2 diabetes: toxic islet amyloid polypeptide oligomers form intracellularly in the secretory pathway. Am J Pathol 176:861–869. doi:10.2353/ajpath.2010.090532

15. Wong WPS, Scott DW, Chuang C-L, et al (2008) Spontaneous diabetes in hemizygous human amylin transgenic mice that developed neither islet amyloid nor peripheral insulin resistance. Diabetes 57:2737–2744. doi:10.2337/db06-1755

16. Mosmann T (1983) Rapid colorimetric assay for cellular growth and survival: application to proliferation and cytotoxicity assays. J Immunol Methods 65:55–63.

17. Trikha S, Jeremic AM (2011) Clustering and internalization of toxic amylin oligomers in pancreatic cells require plasma membrane cholesterol. J Biol Chem 286:36086–36097. doi:10.1074/jbc.M111.240762

18. Costes S, Langen R, Gurlo T, et al (2013) β-Cell failure in type 2 diabetes: a case of asking too much of too few? Diabetes 62:327–335. doi:10.2337/db12-1326

19. Glabe CG (2008) Structural Classification of Toxic Amyloid Oligomers. J Biol Chem 283:29639–29643. doi:10.1074/jbc.R800016200

20. Landreh M, Sawaya MR, Hipp MS, et al (2016) The formation, function and regulation of amyloids: insights from structural biology. J Intern Med 280:164–176. doi:10.1111/joim.12500

21. Zhao H-L, Sui Y, Guan J, et al (2009) Amyloid oligomers in diabetic and nondiabetic human pancreas. Transl Res J Lab Clin Med 153:24–32. doi:10.1016/j.trsl.2008.10.009

22. Lima LMTR (2017) Prediabetes definitions and clinical outcomes. Lancet Diabetes Endocrinol 5:92–93. doi:10.1016/S2213-8587(17)30011-6

23. Lima LMTR (2017) Subclinical Diabetes. An Acad Bras Ciênc. doi: http://dx.doi.org/10.1590/0001-3765201720160394

24. Lin C-Y, Gurlo T, Kayed R, et al (2007) Toxic human islet amyloid polypeptide (h-IAPP) oligomers are intracellular, and vaccination to induce anti-toxic oligomer antibodies does not prevent h-IAPP-induced beta-cell apoptosis in h-IAPP transgenic mice. Diabetes 56:1324–1332. doi:10.2337/db06-1579

25. Zhang S, Liu H, Chuang CL, et al (2014) The pathogenic mechanism of diabetes varies with the degree of overexpression and oligomerization of human amylin in the pancreatic islet β cells. FASEB J Off Publ Fed Am Soc Exp Biol 28:5083–5096. doi:10.1096/fj.14-251744

26. Ashburn TT, Han H, McGuinness BF, Lansbury PT (1996) Amyloid probes based on Congo Red distinguish between fibrils comprising different peptides. Chem Biol 3:351–358.

27. Younan ND, Viles JH (2015) A Comparison of Three Fluorophores for the Detection of Amyloid Fibers and Prefibrillar Oligomeric Assemblies. ThT (Thioflavin T); ANS (1-Anilinonaphthalene-8-sulfonic Acid); and bisANS (4,4'-Dianilino-1,1'-binaphthyl-5,5'-disulfonic Acid). Biochemistry (Mosc) 54:4297–4306. doi:10.1021/acs.biochem.5b00309

28. Kayed R, Head E, Thompson JL, et al (2003) Common structure of soluble amyloid oligomers implies common mechanism of pathogenesis. Science 300:486–489. doi:10.1126/science.1079469

29. Zraika S, Hull RL, Verchere CB, et al (2010) Toxic oligomers and islet beta cell death: guilty by association or convicted by circumstantial evidence? Diabetologia 53:1046–1056. doi:10.1007/S00125-010-1671-6

